# FGF21 acts in the brain to drive macronutrient-specific changes in behavioral motivation and brain reward signaling

**DOI:** 10.1101/2024.03.05.583399

**Authors:** Md Shahjalal H. Khan, Sora Q. Kim, Robert C. Ross, Florina Corpodean, Redin A. Spann, Diana A. Albarado, Sun O. Fernandez-Kim, Blaise Clarke, Hans-Rudolf Berthoud, Heike Münzberg, David H. McDougal, Yanlin He, Sangho Yu, Vance L. Albaugh, Paul Soto, Christopher D. Morrison

## Abstract

Dietary protein restriction induces adaptive changes in food preference, increasing protein consumption over carbohydrates or fat. We investigated whether motivation and reward signaling underpin these preferences. In an operant task, protein-restricted male mice responded more for liquid protein rewards, but not carbohydrate, fat, or sweet rewards compared to non-restricted mice. The protein restriction-induced increase in operant responding for protein was absent in *Fgf21*-KO mice and mice with neuron-specific deletion of the FGF21 co-receptor beta-Klotho (*Klb*^*Cam2ka*^) mice. Fiber photometry recording of VTA dopamine neurons revealed that oral delivery of maltodextrin triggered a larger activation as compared to casein in control-fed mice, whereas casein triggered a larger activation in protein-restricted mice. This restriction-induced shift in nutrient-specific VTA dopamine signaling was lost in *Fgf21*-KO mice. These data strongly suggest that the increased FGF21 during protein restriction acts in the brain to induce a protein-specific appetite by specifically enhancing the reward value of protein-containing foods and the motivation to consume them.

## Introduction

It is well established that animals monitor their nutritional state, detecting and adaptively altering metabolism and feeding behavior in response to the restriction of energy, water, or sodium (1-7). These adaptive responses generally involve neural mechanisms that lead to increased motivation to procure and consume the missing nutrient. Several groups including our own have recently focused on the possibility that the restriction of dietary protein intake also triggers adaptive responses, and indeed protein-restricted rodents selectively shift nutrient preference such that they increase the consumption of protein relative to carbohydrate or fat (8-11). A handful of studies indicate that this ‘protein appetite’ is also associated with an increased motivation for protein, with protein-restricted rats or hamsters exhibiting increased responding for protein or protein-associated cues in an operant task (12, 13) and increased activation of brain reward pathways in response to protein ingestion (14). Taken together, these data suggest that restricting access to dietary protein, or perhaps essential amino acids, induces a selective motivation for protein.

Here we focus on identifying the underlying mechanism driving this protein-specific shift in preference and motivation. For the past several years, we and others have demonstrated that the endocrine hormone Fibroblast Growth Factor 21 (FGF21) is robustly induced by low protein diets and required for animals to adaptively change food intake and nutrient preference during protein restriction (3, 8, 10, 15-17). We, therefore, tested whether FGF21 is also necessary for increased motivation for protein rewards in response to protein restriction. Our results demonstrate that protein-restricted mice exhibit a nutrient-specific increase in motivation for different protein sources and that this protein-specific increase in motivated behavior is dependent on FGF21 and its ability to signal in the brain. In addition, protein restriction enhances the activity of ventral tegmental area (VTA) dopamine neurons in response to oral protein delivery, and this nutrient-specific shift in dopamine neuron response is also FGF21-dependent. Together, these data suggest that FGF21 acts in the brain during protein restriction to selectively increase the responsivity of dopamine neurons to protein while also increasing the motivation to procure protein.

## Results

### Protein-restricted mice exhibit a protein-specific increase in motivated behavior

Our prior work demonstrates that protein-restricted mice shift macronutrient preference, increasing the consumption of protein and reducing the consumption of carbohydrate in a two-choice test (8). Prior work also indicates that protein restriction increases operant responding for protein rewards in rats, consistent with an increase in the motivation for protein (12). We first tested whether this increase in protein motivation would translate to mice, and whether this increased motivation is protein-specific. Male C57BL/6J mice trained to nose poke for liquid rewards were transitioned from standard chow to a low protein (5% casein, LP) or isocaloric control (18% casein, Control) diet and offered various liquid rewards that shifted between casein, maltodextrin, saccharin, sucrose, and corn oil. Mice on LP responded more strongly than control-fed mice when casein was the reward (P = 0.03; **Figure 1**). However, there was no difference in responding between Control and LP mice when any other nutrient served as the reward. Responding at the inactive nose poke was very low and did not differ between groups or across liquid rewards, indicating that active nose poke responding was controlled by liquid reward delivery and was not a random activity. In addition, the LP vs. Control difference in responding for casein was relatively small in the first offering (<5 days of LP diet), but was much larger on subsequent exposures, presumably due to either the increased length of time on the LP diet or familiarity due to repeated exposures. Taken together, these data suggest that protein restriction specifically increases motivation for protein reward.

**Figure 1:**
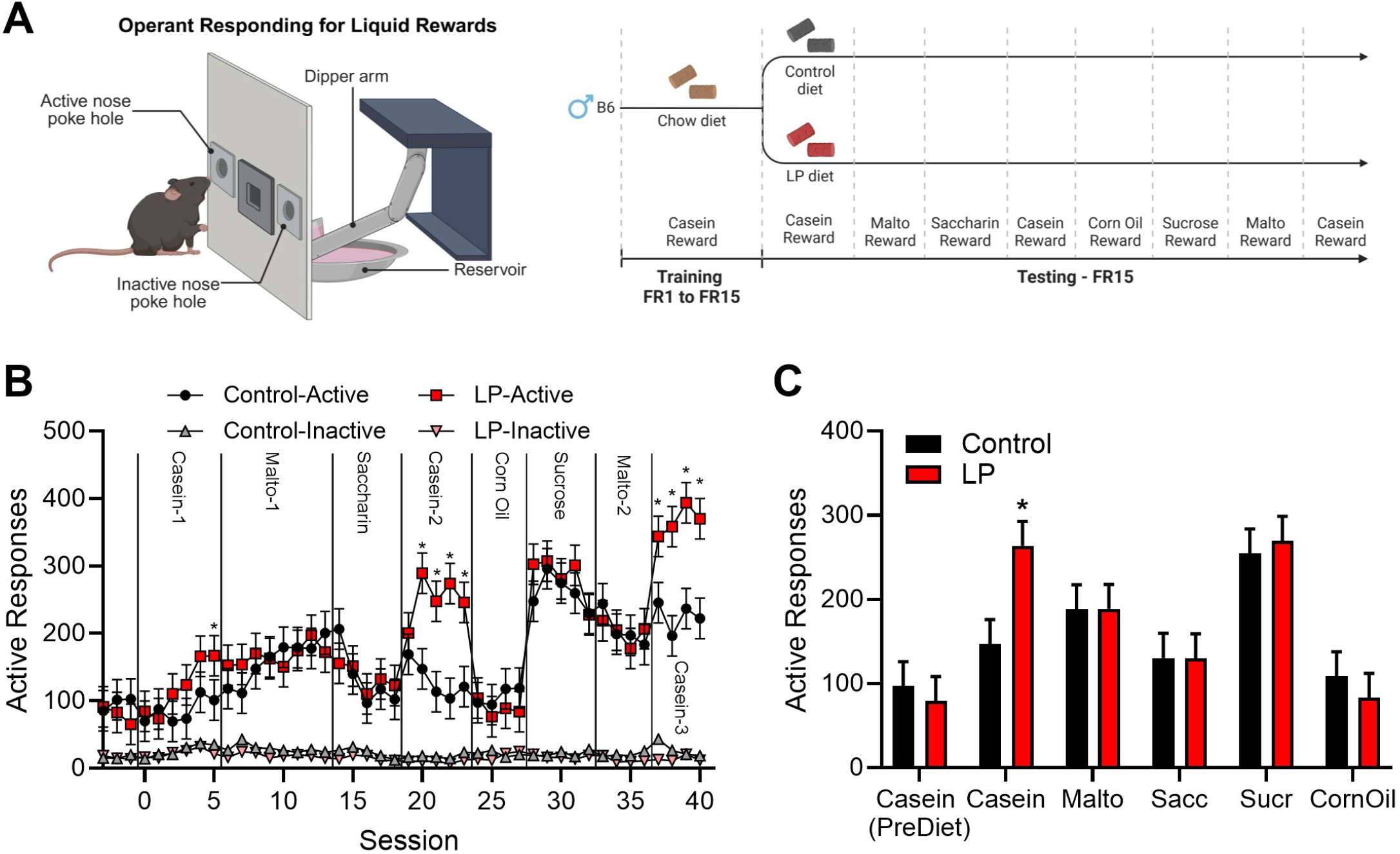
Protein restricted mice selectively increase operant responding for protein. **A**. Male C57BL/6J mice were trained to nose poke for liquid casein rewards under a fixed ratio (FR) 1 schedule and the FR value was subsequently increased to FR15. Half the mice were then transitioned to a low protein (LP) diet and continued to respond for casein before being offered a variety of liquid nutrient rewards. LP mice significantly increased responding for casein, but not for any other nutrient reward **B**. Responses on active and inactive nose poke. **C**. Average active responses for each nutrient. *P<0.05 vs respective control; 8 mice/diet

### Neuronal FGF21 action is required for LP-induced motivation for casein

To test whether FGF21 is required for this LP-induced increase in motivation for protein in male mice, we assessed the demand for protein by progressing control and LP-fed mice through an increasing sequence of fixed ratios: FR1, FR5, FR15, FR45, and FR90 (**Figure 2A**). Consistent with the data above, responding for casein was higher in WT mice on the LP diet compared to WT mice on the control diet at FR values 15, 45, and 90 (**Figure 2B**; Diet x FR P = 0.02). In contrast, responding was not significantly increased by LP diet in *Fgf21*-KO mice at any FR value (**Figure 2C**; Diet x FR P = 0.42). In behavioral economic terms, LP diet-fed mice exhibited greater defense of casein consumption than control diet-fed mice. Following the demand assessment, responding for casein was tested in a PR task (**Figure 2D**). In WT mice, the LP diet tended to increase breakpoint (**Figure 2E**; P = 0.084) and active responses (**Figure 2F**; P = 0.071) during the PR test. In contrast, there was no evidence of an increase in these endpoints in *Fgf21*-KO mice on the LP diet. Taken together, these data suggest that FGF21 is required for LP-induced increases in motivation for protein.

**Figure 2:**
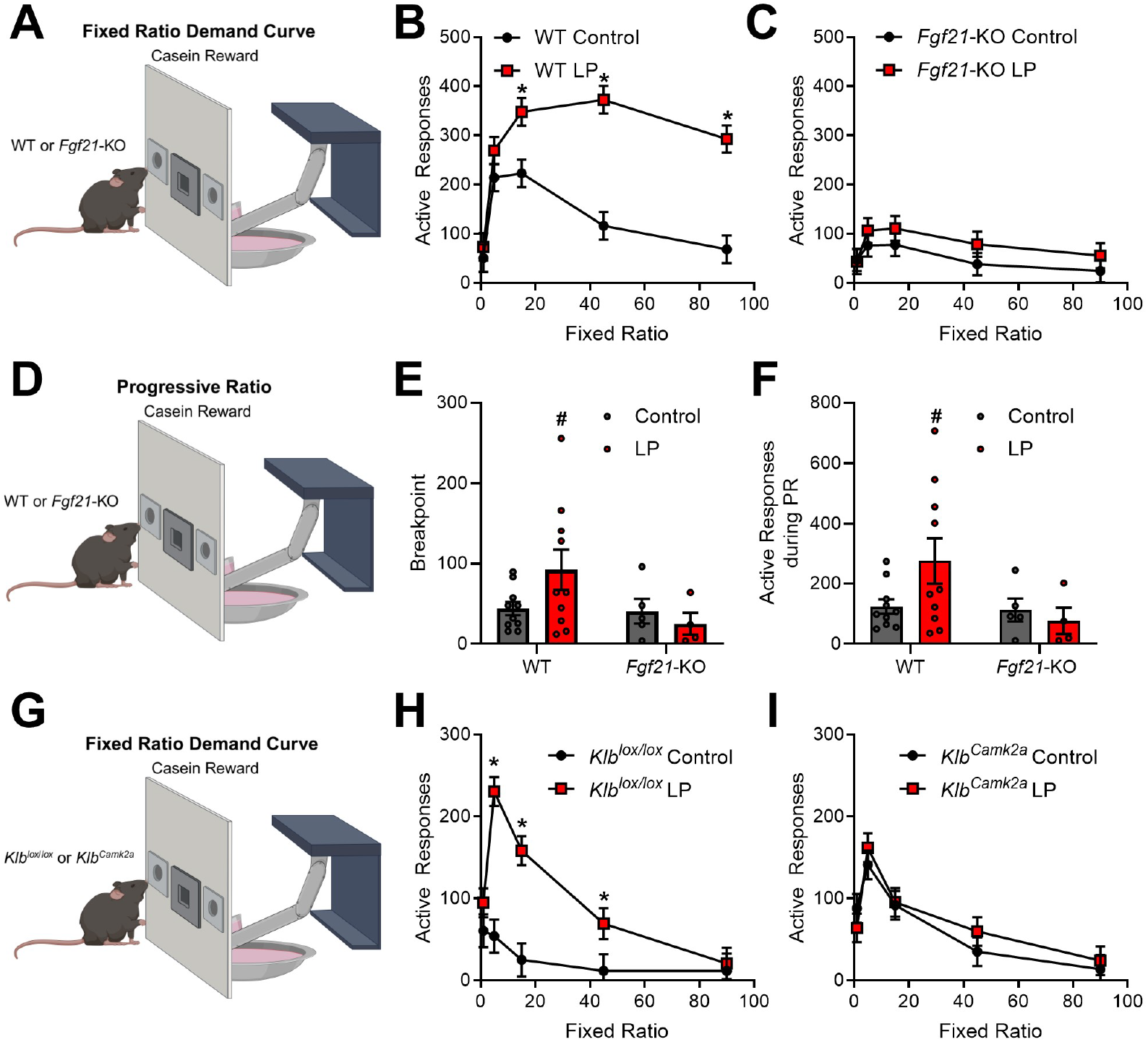
FGF21 signaling in the brain is required for LP-induced increases in operant responding for protein. **A**. Male C57BL6 and Fgf21-KO mice were placed on a control or low protein (LP) diet and trained to nosepoke for casein. **B**. Operant responding for casein in a fixed ratio demand curve in C57BL6 mice (WT; 10 mice/diet). C. Operant responding for casein in a fixed ratio demand curve in Fgf21-KO mice (KO; 4-5 mice/diet). D.Male C57BL6 and Fgf21-KO mice were placed on a control or LP diet and trained to nosepoke for casein. E. Progressive ratio breakpoint in WT and Fgf21-KO mice. **F**. Total active responses during a progressive ratio task in WT and Fgf21-KO mice. G. Male Klb/oxllox and Klb/oxllox;Cam2ka-Cre (Klb^camk*2*a^) were placed on a control or LP diet and trained to nosepoke for casein. **H**. Operant responding for casein in a fixed ratio demand curve in K/b-floxed control (6-8 mice/diet). I. Operant responding for casein in a fixed ratio demand curve in brain-specific Klb knockout mice (Klb^camk*2*a^; 8 mice/diet).*P<0.05; #p< 0.10 vs respective control.

Our prior work suggests that LP-induced changes in food intake and macronutrient preference are largely driven by FGF21 signaling directly within the brain (8). To test whether LP-induced protein motivation also requires brain FGF21 action, mice bearing neuron-specific deletion of Klb (*Klb*^*Camk2a*^) or their floxed littermate controls (*Klb*^*lox/lox*^) were tested using the same demand assessment described above (**Figure 2G**). In *Klb*^*lox/lox*^ mice, the LP diet again increased responding for casein (**Figure 2H**, Diet x FR P= 0.001). However, there was no effect of LP diet on responding in *Klb*^*Camk2a*^ mice (**Figure 2I**; Diet x FR P = 0.73). Taken together, these data indicate that LP-induced increases in motivation for casein require FGF21 signaling in the brain.

### Protein restriction increases the preference and motivation for whey protein

All of the work described above utilized casein as the protein source, and thus the observed behaviors may be driven by sensory properties that are unique to casein. We therefore tested whether LP-induced changes in motivation translate to other protein sources, using whey as the protein source. We first used a 24-hr two-bottle choice model to test the consumption of 4% whey solution vs. 4% maltodextrin (**Figure 3A**). Comparing male WT vs *Fgf21*-KO mice, we observed a significant interaction between diet and genotype for whey consumption, total liquid consumption, and whey preference (**Figure 3B. 3C**; all Ps < 0.01), with WT mice on LP diet consuming more whey (P < 0.001) and showing higher preference (P = 0.0015) relative to control fed mice. In contrast, LP did not increase whey intake and preference in *Fgf21*-KO mice.

**Figure 3:**
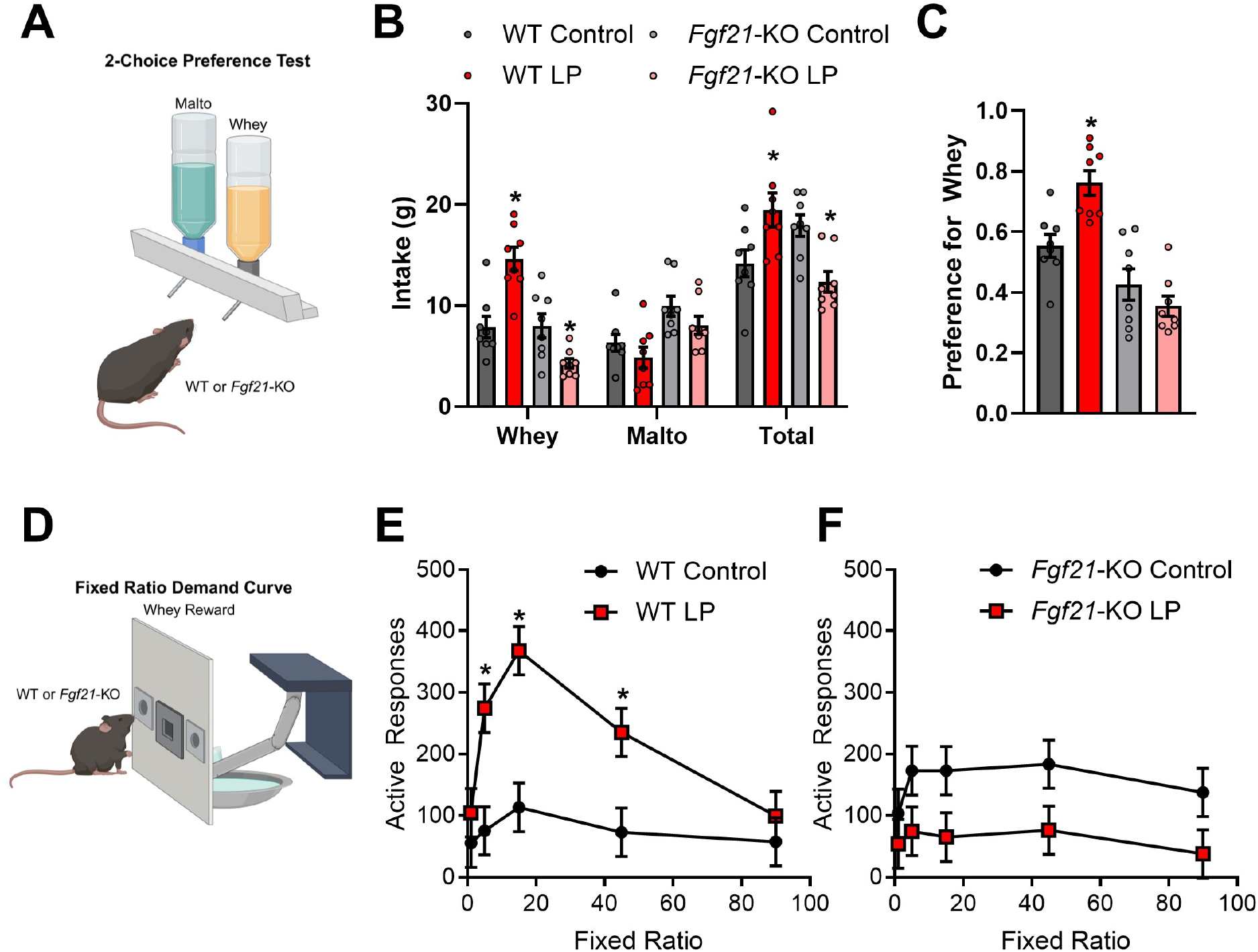
Protein restriction increases preference and motivation for whey protein in WT but not *Fgf21-KO* mice. **A**. Male mice were placed on a control or low protein (LP) diet for 7 days, and were then offered the choice beween 4% whey and 4% maltodextrin solutions in the home cage over three 24-hr periods. **B**. Average 24hr consumption of whey and malto. **C**. Preference ratio for whey (whey consumption divided by total consumption). **D**. Male mice on control or LP diet were trained to nosepoke for whey in a rapid demand curve assessment. **E**. Operant responding for whey in a fixed ratio demand curve in C57BL/6J mice (WT). **F**. Operant responding for whey in a fixed ratio demand curve in *Fgf21-KO* mice (KO). *P<0.05 vs respective control; 8 mice/group)

We then moved to an operant responding paradigm similar to that described above, except that after mice were trained to respond at FR1, only one operant session occurred at each FR value (**Figure 3D**). In this more rapid protocol, we observed a significant interaction between diet and genotype (P = 0.045), with WT mice on LP diet responding more than control mice for whey (**Figure 3E**) but this effect being completely lost in *Fgf21*-KO mice who reduced responding for whey when on LP diet (**Figure 3F**). Taken together, these data reinforce the concept that protein restriction significantly increases the preference for protein relative to carbohydrate as well as the motivation to procure protein. This LP-induced effect manifests across multiple protein sources and requires FGF21 signaling.

### FGF21 is required for nutrient-specific shifts in VTA dopamine neuron activity in protein-restricted animals

Dopamine neurons within the VTA are closely linked to reward and motivation, and recent work in rats suggests that protein restriction enhances protein-induced activation of unidentified VTA neurons (14). We, therefore, sought to use mice to specifically test whether dopamine neurons within the VTA respond to the delivery of protein within the oral cavity, and if this response is enhanced by protein restriction. To test this question, *Th*-Cre mice were used for fiber photometry recording of VTA neurons in response to the intraoral delivery of protein (4% casein) or carbohydrate (4% maltodextrin) (**Figure 4A**). We tested the relative strength of casein vs. maltodextrin-induced neural activity to assess macronutrient-specific differences as compared to broader nutrient-agnostic effects. When male WT mice consuming the control diet were tested, intraoral delivery of maltodextrin or casein both produced distinct increases in VTA dopamine neuron activity (**Figure 4B, 4C**). However, the malto-induced activation was relatively larger than the casein-induced activation, with maltodextrin inducing a larger mean signal (P = 0.055, **Figure 4D**), but not the maximum (peak) signal (**Figure 4E**), relative to casein in the 20 seconds post-infusion.

**Figure 4:**
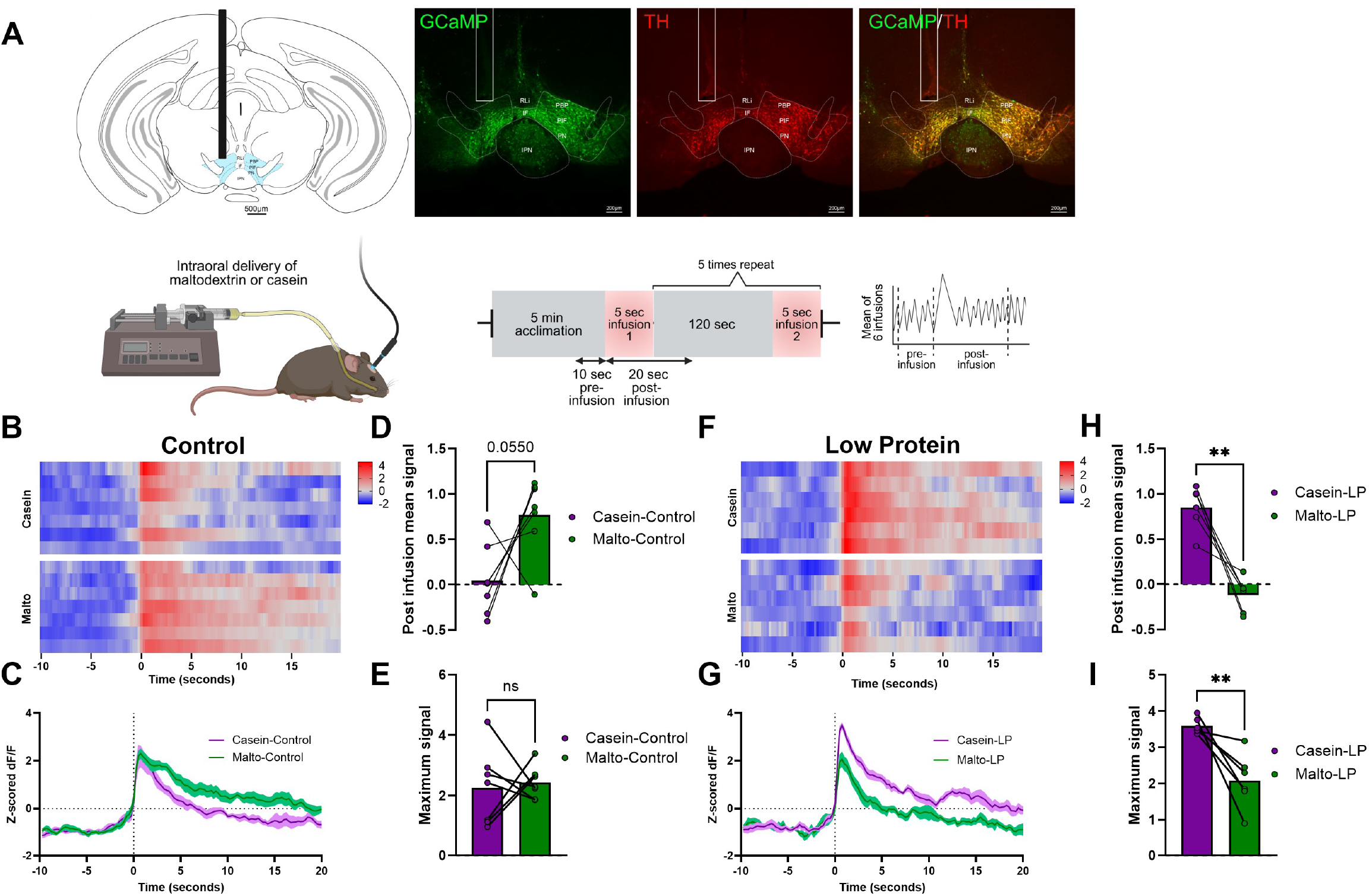
Protein restriction induces a macronutrient-dependent shift in VTA-dopamine neuron activity. **A**.Fiber photometry recording of TH-positive neurons in the VTA was conducted in male mice on control diet in response to intraoral delivery of either 4% casein or 4% maltodextrin solutions. Mice were then placed on LP diet for 1 0 days and the recording was repeated. B. Heat map of neuron activity in individual animals on Control diet, with each row representing z-scored dF/F signal from a single animal. C. Average change in neuronal activity in response to nutrient infusion in Control-fed animals. Solid line indicates mean and shaded area represents SEM. **D**. Mean signal during the 20 seconds post intraoral infusion on control diet. **E**. Maximum signal during the 20 seconds post intraoral infusion on control diet. **F**. Heat map of neuron activity in individual animals on LP diet, with each row representing z-scored dF/F signal from a single animal. **G**. Average change in neuronal activity in LP-fed animals. Solid line indicates mean and shaded area represents SEM. **H**. Mean signal during the 20 seconds post intraoral infusion on LP diet. I. Maximum signal during the 20 seconds post intraoral infusion on LP diet. *P<0.05, **P<0.01 vs. respective control; 6 mice/group. Modified schematic of mouse brain atlas adapted from Paxinos and Franklin (53).

Interestingly, these macronutrient-induced neuron responses were reversed when mice were fed the LP diet. In this case, casein induced a larger increase compared to maltodextrin (**Figure 4F. 4G)**, resulting in a larger mean (**Figure 4H;** P < 0.01) and maximum (**Figure 4I;** P < 0.01) signal compared to maltodextrin in the 20 seconds post intraoral infusion. Collectively, these data indicate that LP shifts the relative strength of the VTA dopamine neuron response, increasing the size of the casein vs maltodextrin-induced dopamine response.

Considering FGF21’s role as a signal of the protein-restricted state, we then tested whether this LP-induced shift in VTA dopamine neuron activity requires FGF21. To address this question, the *Th*-Cre allele was crossed onto the *Fgf21*-KO background, and these *Th*-Cre;*Fgf21*-KO mice were used to test nutrient-induced activation of VTA dopamine neurons via fiber photometry as described above (**Figure 5A**). In the absence of FGF21, we again observed readily apparent increases in VTA dopamine neuron activity in response to the oral delivery of either casein or maltodextrin. Maltodextrin produced a relatively larger dopamine activation as compared to casein (**Figure 5B. 5C)**, with a significantly larger mean signal in the 20-second post-infusion period (**Figure 5D**; P < 0.01). However, the LP-induced shift in the relative strength of these responses was absent in *Th*-Cre;*Fgf21*-KO mice, such that maltodextrin continued to produce a relatively larger increase in dopamine neuron activity when mice consumed LP diet (**Figure 5G. 5H, 5I**, P < 0.05). Taken together, these data suggest that FGF21 is essential for macronutrient-specific shifts in dopamine neuron activity in protein-restricted animals.

**Figure 5:**
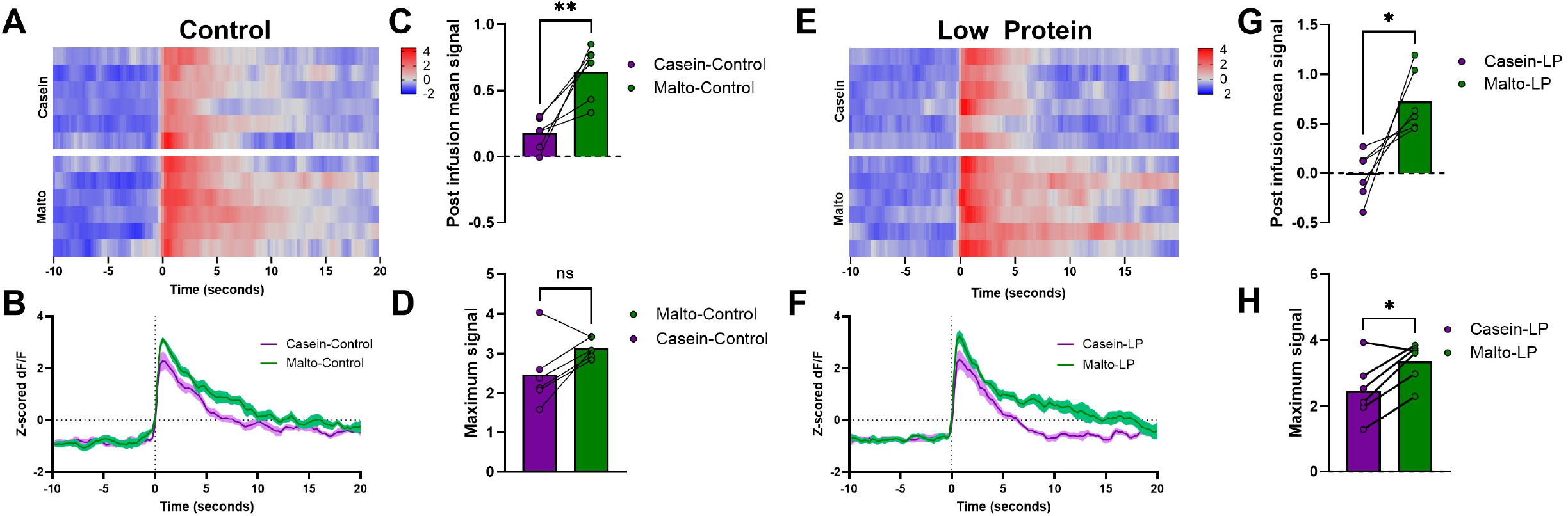
Deletion of FGF21 blocks the effects of protein restriction on VTA-dopamine neuron activity. Male *Th-Cre;Fgf21-KO* mice on Control diet were used to assess fiber photometry recording of TH-positive neurons in the VTA in response to intraoral delivery of either 4% casein or 4% maltodextrin solutions. Mice were then placed on LP diet for 10 days and the recording was repeated. A. Heat map of neuron activity in individual animals, with each row representing z-scored dF/F signal from a single animal. **B**. Average change in neuronal activity on control diet. Solid line indicates mean and shaded area represents SEM. **C**. Mean signal during the 20 seconds post intraoral infusion on control diet. **D**. Maximum signal during the 20 seconds post intraoral infusion on control diet. **E**. Heat map of neuron activity in individual animals, with each row representing z-scored dF/F signal from a single animal. **F**. Average change in neuronal activity on LP diet. Solid line indicates mean and shaded area represents SEM. **G**. Mean signal during the 20 seconds post intraoral infusion on LP diet. H. Maximum signal during the 20 seconds post intraoral infusion on LP diet. *P<0.05, **P<0.01 vs. respective control; 6 mice/group

## Discussion

Prior work has established that protein restriction alters macronutrient intake (9, 18, 19). Protein-restricted mice increase total food intake if maintained exclusively on a low protein diet, but selectively shift their preference towards protein-rich foods if given a choice between various nutritional options (8-11, 15). In addition, these changes in protein intake require FGF21 signaling in the brain, as the deletion of either FGF21 from the whole animal or deletion of its receptor in neurons blocks these adaptive changes in food intake in response to protein restriction (8, 15, 20). Here, we focus on the mechanisms that drive these FGF21-dependent changes in macronutrient preference, focusing specifically on motivation, reward, and dopamine signaling.

Motivation drives food intake in various need states, and therefore one would predict that a protein-restricted state would increase the value of protein and lead animals to work harder to attain protein-rich rewards (21). Although there has been very little work related to protein-specific motivation, prior work has shown that protein restriction increases operant responding for protein in both golden hamsters (13) and rats (12). Therefore, we sought to extend these data to mice, enabling the use of mouse genetics to test the role of FGF21 in driving these changes in motivation during protein restriction.

We first sought to establish a model to assess motivation for liquid rewards in mice, as liquid rewards provide a means to easily test various nutrients in discrete amounts over time. We observed clear and significant increases in operant responding for casein in LP-fed mice as compared to control mice, but interestingly there was no LP effect for any other nutrient tested, including maltodextrin, sucrose, corn oil, and saccharin. These results provide strong evidence that the LP-induced increase in motivation is nutrient-specific and that the hyperphagia observed in animals maintained exclusively on a low protein diet is driven by a specific motivation to consume protein (17). Indeed, our prior data indicate that protein-restricted mice do not increase maltodextrin consumption when it is offered alone, nor do they consistently increase total food intake if they can choose between high and low protein diets (8). These observations are consistent with the concept of protein leveraging, which predicts that animals prioritize protein over energy if they cannot select between foods to balance their energy vs. protein needs (17, 22, 23).

Interestingly, we did not observe any difference in operant responding for sucrose, despite evidence that FGF21 inhibits sweet intake (24, 25). Based on this prior work and the fact that the LP diet induces a large increase in FGF21 (8, 15, 20), we anticipated that mice on LP might reduce operant responding relative to control mice when sucrose or saccharine was the reward. It seems possible that LP-induced FGF21 signaling drives different mechanisms than exogenous FGF21 administration, or that other pathways activated in the low protein state serve to blunt the effects of FGF21 on sweet intake. Interestingly, recent work indicates that a low-protein meal does not alter the consumption of a sweet dessert but does increase the consumption of a high-protein dessert (26), while the well-established effects of carbohydrate meals to drive FGF21 were blocked by relatively small amounts of added protein (27). Recent work in preprint indicates that protein-restricted mice exhibit decreased consumption for sucrose, decreased motivation for sucrose in a conditioned place preference task, and decreased dopamine release in the nucleus accumbens (28). It is unclear why protein-restricted mice did not show a reduction in operant responding for sucrose in this study, and more focused work on protein restriction and its effects on motivation for sweet rewards would be necessary to resolve this question.

Having established a model in which protein-restricted mice exhibit a clear increase in motivation for protein, we next focused on the mechanisms that might drive this protein-specific motivation. As described above, our prior work indicates that FGF21 is essential for LP-induced hyperphagia in mice on single-choice diets (15, 20), as well as LP-induced changes in protein vs carbohydrate preference in two-choice models (8). We, therefore, tested whether FGF21 is also required for protein restriction to increase motivation for casein, using a fixed ratio demand curve assessment in which the effort to procure casein increased with time. We observed that the LP diet markedly increased operant responding for casein in WT mice, suggesting an increase in motivation for protein. In contrast, this LP effect was completely lost in *Fgf21*-KO mice. This demand curve approach was replicated in a different line of mice bearing brain-specific deletion of the FGF21 co-receptor Klb (*Klb*^*Camk2a-Cre*^), and again we observed that the LP diet increased operant responding for casein in the floxed controls, while this LP-effect was completely lost in the brain-specific Klb knockouts. These data, generated from two separate genetic lines, support the overarching hypothesis that FGF21 acts in the brain to promote protein intake by inducing a protein-specific motivation.

LP-induced changes in casein consumption or motivation might be driven by some property inherent to casein that is unrelated to its nutritional value. To confirm that protein restriction broadly increased consumption and motivation for protein, we replicated our core observations using whey, since whey protein contains a different amino acid composition and digestibility profile compared to casein (29-31). We first tested whether protein restriction altered whey consumption or preference in a 2-choice paradigm with maltodextrin. Just as with casein (8, 11), mice on LP diet significantly increased their consumption and preference for whey vs. maltodextrin. Importantly, this shift in macronutrient preference was lost in *Fgf21*-KO mice, again demonstrating FGF21’s importance in mediating these behavioral responses. We then tested whether protein restriction altered motivation for whey, using a rapid demand curve operant paradigm. Here again, mice on LP diet increased responding for whey in a manner analogous to mice responding for casein, and again this LP-induced increase in responding was lost in *Fgf21*-KO mice. Taken together with the operant data in Figures 1 and 2. these data strongly indicate that protein restriction induces a broad increase in consumption, preference, and motivation for protein foods and/or protein rewards, but that these effects do not extend to other macronutrients. As such, these data suggest that protein-restricted mice manifest a ‘protein appetite,’ with protein restriction causing animals to specifically seek and consume protein without increasing the consumption of other foods. The changes in feeding behavior driven by protein restriction are fundamentally different from the effects of energy restriction or general dietary (food) restriction, which promote hyperphagia and increased motivation to procure food in general. Finally, and importantly, these data again support the essential role of FGF21 in mediating these protein-specific changes in motivation and consumption.

Because these data demonstrate that protein restriction leads to an FGF21-dependent increase in the rewarding value of protein, we next focused on the neural mechanisms mediating this behavior. The homeostatic detection and defense against nutrient restriction is mediated by multiple brain areas, including those classically associated with reward (32, 33). The mesolimbic dopamine system is particularly linked with reward behavior, including the response to food and food rewards (34-38). Indeed, neurons in the VTA were activated by protein intake in rats, and this activation was enhanced by protein restriction (14). Similarly, meal-induced cFos in the nucleus accumbens was enhanced in protein-restricted rats consuming a high-protein meal (39). We, therefore, tested whether we could replicate this effect in mice, as mouse genetics allows the specific targeting of VTA dopamine neurons (via *Th*-Cre) and testing their activity in the absence of FGF21 (via *Fgf21*-KO mice). Delivery of either casein or maltodextrin into the oral cavity produced a clear, temporally discrete increase in VTA-dopamine neuron activity. In mice consuming a control diet, the relative response of VTA dopamine neurons to intraoral maltodextrin was slightly but consistently larger than the response to intraoral casein. These effects are consistent with our observation that control-fed mice prefer maltodextrin to casein in a 2-choice preference test (8). Conversely, in mice consuming the LP diet, casein produced a larger activation of dopamine neurons compared to maltodextrin, again consistent with the fact that LP-fed mice prefer casein. Thus, these data suggest that protein restriction induces macronutrient-specific shifts in the response of the reward system to orally delivered nutrients. Finally, we point out that this comparison of casein to maltodextrin-induced signal is critical for disentangling macronutrient-specific effects from broad, nutrient-independent changes. Both casein and maltodextrin induce VTA dopamine signaling and are readily consumed, regardless of whether the animal is on the control or LP diet. However, protein-restricted mice exhibit a selective shift in the relative value of these two macronutrients, such that carbohydrate is more valued in control animals and protein is more valued in protein-restricted animals. These behavioral changes are mirrored by changes in the relative strength of dopamine neuron activation following oral delivery.

Finally, we tested whether these diet-induced shifts in macronutrient-induced dopamine neuron activity were dependent on FGF21, using *Th*-Cre;*Fgf21*-KO to selectively record from VTA dopamine neurons in the absence of FGF21. We again observed that VTA dopamine activity was increased by the intraoral infusion of both maltodextrin and casein, but the strength of the casein vs. maltodextrin-induced signal did not shift when *Th*-Cre;*Fgf21*-KO mice were fed LP. As such, not only do *Fgf21*-KO mice not change their preference or motivation for protein on a LP diet, but the dopamine neuron response to specific macronutrients also does not shift. Therefore, these data are consistent with the work above and collectively support a model in which FGF21 acts in the brain to promote protein intake by altering the reward system response to nutrient ingestion in a manner that enhances the value of protein.

While our work broadly supports a model in which FGF21 acts in the brain to increase protein motivation, there are several caveats to consider with this work. First, where and how FGF21 acts to alter neural activity in the VTA is unclear. Available evidence suggests that Klb is not heavily expressed within the VTA (40, 41), and thus it is likely that these effects are mediated indirectly in response to FGF21 action in upstream brain areas. Second, although it is well established that VTA dopamine neurons play a critical role in motivation and learning, our data do not definitively demonstrate that changes in dopamine neuron activity drive the changes in operant responding that we observe during protein restriction. Indeed, it is somewhat simplistic to view dopamine neuron activation as strictly encoding reward, as dopamine neurons respond to a variety of stimuli, including aversive stimuli. Finally, despite specific expression of TH in dopamine neurons (42), we acknowledge that this TH-Cre strategy inevitably targets a small number of non-dopaminergic neurons (43, 44). However, this non-specificity is primarily detected in areas medial (RLi and IF) or outside the VTA (IPN), which are not targeted in our study. Instead, our fiber optic cannula targets the lateral VTA (PBP, PIF, PN), where there is a high degree of specificity in the TH-Cre model. While no animal model is perfect, we are confident that our work primarily measures dopamine neuron activity in the lateral VTA.

In conclusion, our data provide strong evidence that protein restriction induces a macronutrient-specific increase in motivation for protein. Protein-restricted mice will work harder for multiple protein sources in an operant task, but they do not work harder for carbohydrate, sweet, or fat. In addition, these behavioral changes are accompanied by changes in the mesolimbic reward system, with protein restriction altering the dopamine neuron response to nutrient ingestion in a macronutrient-specific fashion. Finally, both the behavioral and cellular manifestations of protein reward are dependent on FGF21, and most likely its ability to signal in the brain. As such, these data provide convincing evidence that FGF21 is an endocrine signal of protein restriction that acts in the brain to specifically enhance the reward value of protein-containing foods and promote their consumption. FGF21 thus provides a compelling mechanism to explain how animals within complex nutritional landscapes effectively balance the physiological need for protein vs. other macro and micronutrients.

## Acknowledgments

The authors would like to thank the leadership and staff of the PBRC Comparative Biology Core and Animal Metabolism and Behavior Core for their skillful assistance and excellent technical support. This work was supported by the National Institutes of Health (NIH) R01DK123083 and S10OD023703 to C.D.M. SQK was supported by T32DK064584. RAS was supported by F32DK130544. This project used facilities within the Animal Metabolism & Behavior Core, Genomics Core, and Cell Biology and Bioimaging Core at PBRC that are supported in part by NIH center awards P20GM135002 and P30DK072476, as well as an NIH equipment award S10OD023703.

## Author Contributions

Conceptualization – MSK, SQK, PS, CDM; Methodology - MSK, SQK, RCR, FC, HRB, YH, PS, CDM; Investigation - MSK, SQK, RCR, FC, RAS, DAA, SFK, BC, SY, VLA, PS; Visualization - MSK, SQK, HRB, VLA, PS, CDM; Formal - analysis - MSK, SQK, PS, CDM; Writing - Original Draft - MSK, SQK, RAS, PS, CDM; Writing - Review & Editing - MSK, SQK, HRB, HM, DM, YH, SY, VLA, PS, CDM; Funding acquisition – CDM.

**The authors declare no competing interests**

## Methods

### Animals and diets

All animal-related procedures were approved by the PBRC Institutional Animal Care and Use Committee (IACUC) and were carried out in strict adherence to the guidelines and regulations set by the NIH Office of Laboratory Animal Welfare. Male C57BL/6J (WT) mice obtained from Jackson Laboratory were used in all experiments. *Fgf21*-KO mice on the B6 background were provided by Dr. Steven Kliewer (45) and bred in the homozygous state with C57BL/6J mice used as controls. Beta-Klotho floxed (*Klb*^*lox/lox*^) mice were provided by Dr. Steven Kliewer (40, 46) and crossed with *Camk2a*-Cre (47) to generate brain-specific Klb knockouts (*Klb*^*Cam2ka*^). *Th*-Cre mice (48) were procured from the European Mouse Mutant Archive (B6.129X1-Th^tm1(cre)Te^/Kieg, EMMA #: EM_00254). *Th*-Cre was bred onto the *Fgf21*-KO background to allow viral targeting and fiber photometry recording from dopamine neurons in the absence of FGF21.

Mice were individually housed in a controlled environment with regulated temperature and humidity and maintained on a 12:12-hr light-dark cycle. Throughout the experiments, the mice had ad-libitum access to food and water unless stated otherwise. All experiments were conducted during the light cycle. The Control and low protein (LP) diets were formulated and produced by Research Diets and were designed to be isocaloric by adjusting the levels of protein and carbohydrates while keeping the fat content constant. The Control diet consisted of 20% casein (by weight) as the primary source of protein, while the LP diet included 5% casein. Diet descriptions have been previously published (8, 20, 49).

### Experiment 1: Protein-restricted mice exhibit nutrient-specific increases in motivated behavior

Experimental sessions were conducted in operant conditioning chambers equipped with two nose poke response devices, a house light for general illumination, and a switchable liquid dipper that provided access to 0.01 ml of an experimenter-made solution. Male C57BL/6J mice (n=16) fed a chow diet were trained to nose poke for a 4% casein/0.1% saccharin solution by raising the dipper for 10 s each time the mouse made a nose poke response and providing response-independent dipper access at variable times with inter-delivery intervals averaging 6 min. Once a mouse made 10 responses within a single session, response-independent solution deliveries were discontinued, and only responses in the “active” nose poke hole (left/right assignment of the active nose poke hole was counterbalanced across mice) produced solution access. Once mice exhibited stable responding under a fixed ratio (FR) 1 schedule of reinforcement, the FR value was increased progressively to 15 (15 nose pokes = 1 reward), and the FR value held constant at 15 for the remainder of the experiment. At this point, mice were randomized to either control or LP diet (8/diet) for the remainder of the study (day 0), and across sessions, the liquid reward was altered between multiple nutrient sources (Figure 1): 4% casein/0.1% saccharin, 4% maltodextrin/0.1% saccharin, 0.1% saccharin, 15% sucrose, and corn oil. Sessions occurred once per day, with mice offered each nutrient reward for 4-8 sessions before a different reward was offered.

### Experiment 2: FGF21 action in the brain is required for LP-induced increases in motivation for protein

Training and experimental sessions were conducted using operant conditioning chambers described in Experiment 1. Male C57BL/6J (WT, n=20) and *Fgf21*-KO (KO, n=9) mice were randomly assigned to either control or LP diet for 10 days and subsequently trained to nose poke for 4% casein/0.1% saccharin solution as described above. Once mice exhibited stable responding at FR 1, a demand assessment was conducted by varying the FR value in ascending order: 1, 5, 15, 45, and 90, with at least 5 individual sessions at each FR value (50). Following the sequence of FR values, mice were tested in two separate progressive ratio sessions (on separate days), in which the number of nose pokes required to elicit a reward progressively increased within the session (51), based on the following formula: Ratio Value = 5*exp(Ratio Number*0.2)-5. Progressive ratio (PR) sessions ended when mice failed to respond after 10 minutes (breakpoint) or at the end of the session.

A separate study was conducted to replicate these findings in a group of brain-specific Klb knockout mice. Male Cre-negative *Klb*^*lox/lox*^ mice or Camk2-Cre positive littermates (*Klb*^*Camk2a*^) were randomized to control or LP diet and trained to nose poke for casein solution as above (6-8 mice/genotype/diet). Mice were then tested over increasing FR values of 1, 5, 15, 45, and 90, as described above.

### Experiment 3: Protein restriction increases preference and motivation for whey in WT but not Fgf21-KO mice

Male C57BL/6J and *Fgf21*-KO (KO) mice were placed on Control or LP diet (8 mice/diet/genotype) for 7 days and were then offered two bottles containing either 4% whey/0.1% saccharin or 4% maltodextrin/0.1% saccharin for 3 days. Bottle locations were counterbalanced across mice and locations were swapped each day. Fluid consumption was measured daily and averaged across the 3-day experiment to provide average daily consumption. Preference for whey, calculated for each animal, was derived from average daily whey consumption divided by total consumption.

A separate group of male C57BL/6J and *Fgf21*-KO (KO) mice were trained to nose poke for 4% whey/0.1% saccharin as described in Experiments 1 and 2 above, except the study was compressed into a shorter time window in which only a single session was conducted at each FR. Mice were placed on either control or LP diet (8 mice/diet/genotype) for 10 days followed by training sessions and initial sessions at FR1 (as described above). When all the mice fulfilled the initial training criteria and stably responded at FR1, mice were tested over an increasing series of FR values of 1, 5, 15, 45, and 90, with only one session conducted at each FR value.

### Experiment 4: FGF21 is required for increased activation of VTA dopamine neurons to protein in protein-restricted animals

A Cre-dependent GCaMP6s virus (AAV-DJ-EF1a-DIO-GCaMP6; Stanford Gene Vector and Virus Core) was stereotaxically delivered to the VTA of male *Th*-Cre mice (n=6), and a fiber optic cannula (Doric Lenses) was implanted at the injection site to record neural activity. Mice were given 3 weeks to recover, at which point fiber photometry recording (RZ10x Fiber Photometry Processor with integrated LUX LEDs and Photosensors; Tucker-Davis Technologies) of VTA dopamine neurons was conducted to confirm active GCaMP signal. Mice were then placed on a control (CON: 20% casein) diet for 10 days, an intraoral cannula was implanted, and mice were acclimated to intraoral water infusions. Mice were also given a single overnight exposure to the solutions (4% casein and 4% maltodextrin) before the initiation of recordings. To record neural activity to casein or maltodextrin, intraoral infusions occurred during 15-minute sessions. After an initial 5-minute acclimation period, each mouse received 6 separate intraoral presentations of 4.8 μl/s for 5 seconds), with each presentation separated by at least 120 seconds. VTA neural activity was recorded continuously throughout the session. Casein and maltodextrin were provided in two different sessions (morning and afternoon) on the same day, ∼5-6hrs apart, with presentation order randomized per animal. After recording under the control diet, all mice were shifted to the LP diet (LP: 5% casein) for another 10 days, and the dopamine neuron activity during intraoral casein or maltodextrin infusion was again recorded as above. For the analysis, custom MATLAB scripts were used to fit the GCaMP signal (465nm) to the isosbestic control signal (405nm), calculating dF/F. For each presentation, the mean baseline signal (10 seconds pre-infusion) was subtracted from each data point across the 30-second experiment period. Then, the baseline-adjusted data were averaged across the 6 presentations per session per animal, generating one curve of intraoral infusion-induced signal spanning 10 seconds pre-infusion and 20 seconds post-infusion. To account for the inter-animal differences in signal intensities, Z-scores were calculated for each mouse using the mean and standard deviation of the 30-second experiment period across casein and maltodextrin test sessions. Using the z-standardized values, the mean and maximal (peak) signal for the 20 seconds after infusion were calculated and compared.

To test whether LP-induced changes in dopamine neuron activity require FGF21, the *Th*-Cre allele was bred onto a homozygous *Fgf21*-KO background. *Th*-Cre;*Fgf21*-KO mice (n=6) were then placed on control and LP diet, with the dopamine neuron response to intraoral delivery of casein and maltodextrin measured as described above.

### Stereotaxic brain surgery and cannulation procedure

Mice were anesthetized using isoflurane (2-5%) and positioned under a stereotaxic instrument. A midline incision was made on the skin, and the skull surface was leveled. The bregma landmark was identified and used to adjust the positioning. A 1 mm hole was drilled in the skull to accommodate the guide cannula (Plastics One), which was then lowered into the brain, precisely above the VTA target. A unilateral delivery of a recombinant Adeno-associated virus (rAAV) solution was performed, and the injection coordinates relative to bregma were as follows: anterior-posterior (AP) -3.3, medial-lateral (ML) +0.4, dorsal-ventral (DV) -4.4 and -4.2 mm for the two injections. Using an injector attached to a 1 μl Hamilton syringe, a total injection volume of 0.8 μl (0.4 μl for each injection) was infused at a rate of 0.1 μl/min. After the first injection, a 5-minute diffusion period was allowed before the cannula was raised to the higher site and the second injection was performed. Following the viral injections, a chronically implantable optic fiber (Doric Lenses, MFC_400/430-0.48_4.3mm_MF2.5_FLT) was implanted at a DV coordinate of -4.3 mm. The optic fiber was secured in place using dental acrylic adhesive cement (C&B-Metabond). Following recovery, all mice were initially tested for signal strength and responsivity of neural activity before inclusion in the study. Following the experiments, mice were euthanized and immunohistochemistry was conducted to confirm appropriate cannula placement (Supplemental Figure 1).

### Intraoral Surgery and cannulation

Intraoral cannulas were placed under general anesthesia (2-5% isoflurane) and subcutaneous analgesia (carprofen) in the animal operating room at Pennington Biomedical Research Center, using a procedure modified from Stratford and Thompson (52). A midline skin incision was made on the dorsal surface immediately caudal to the pinnae. A sterile stainless steel hypodermic tube was inserted through the incision and guided subcutaneously to the oral cavity lateral to the molars and an intraoral incision was made. A 6-cm length of flared polyethylene catheter with a small Teflon washer was inserted intraorally through the hypodermic tube and the tube was removed such that the Teflon washer rested flush against the inner buccal cavity just lateral to the first maxillary molar. The catheter was then secured in place subcutaneously, the skin was closed using non-absorbable suture, and the catheter was flushed with sterile water. Following surgery, mice were provided 0.5ml of warmed sterile saline subcutaneously and mashed food. Once fully recovered, mice were regularly infused with water via the intraoral cannula to ensure patency and acclimate the animal to intraoral fluid delivery.

### Statistical Analysis

Data were analyzed using the SAS version 9 or JMP Pro version 16 software packages (SAS Institute) via one-way, two-way, or repeated-measures ANOVA using the general linear model procedure. When experiment-wide tests were significant, post hoc comparisons were made using the LSMEANS statement with the PDIFF option and represent least significant differences tests for pre-planned comparisons. Fiber photometry data were analyzed as described above, with a two-tailed paired Student’s t-test to assess differences in mean signal and max signal before and after the low protein diet. All data are expressed as mean ± SEM, with a probability value of less than 0.05 considered statistically significant.

### Audio/Visual Credits

All schematics in all figures were Created with BioRender.com

## Notes

### Competing Interest Statement

The authors have declared no competing interest.

### Summary of Updates

Updates to Figure 4 and Discussion

