## Supplemental Figure 1 for "FGF21 acts in the brain to drive macronutrient-specific changes in behavioral motivation and brain reward signaling" (from another version of this preprint: version 1, 2024-03-07)

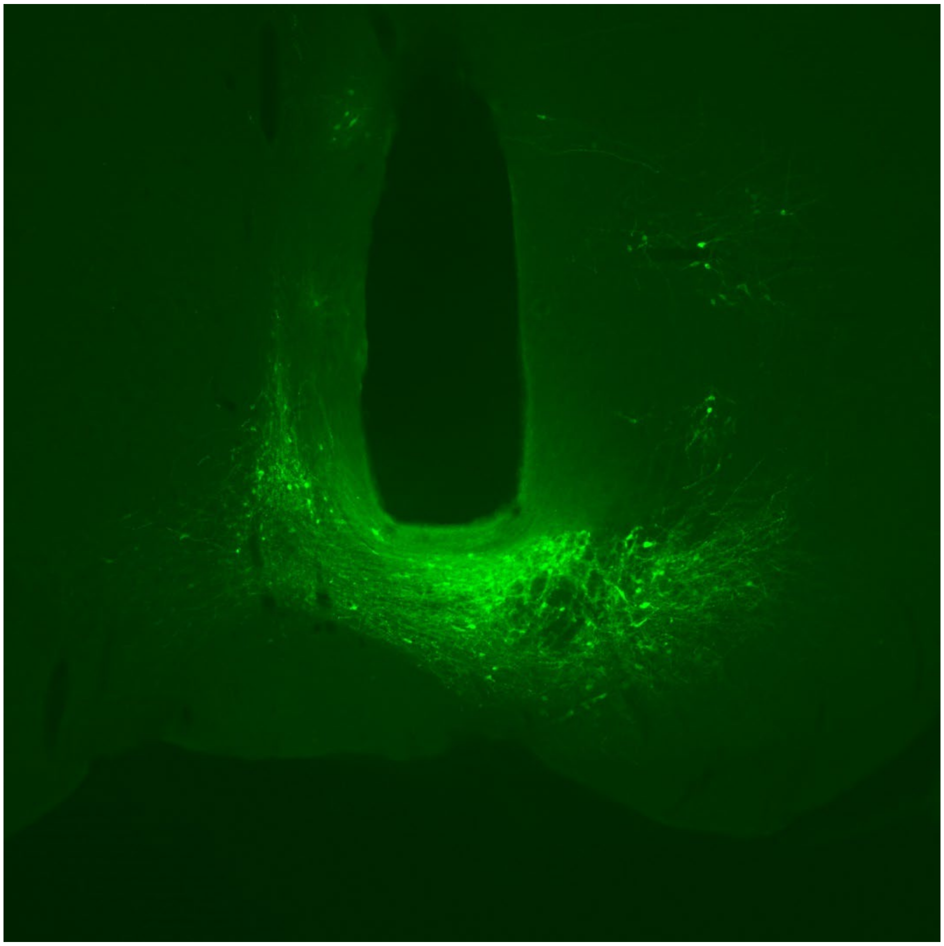

**Supplemental Figure 1: Representative image of AAV-GCaMP targeting the VTA.** *Th*-Cre mice were injected with a Cre-dependent AAV-GCaMP unilaterally into the VTA to target dopamine neurons for fiber photometry recording. After viral injection, and fiberoptic cannula was implanted, targeting the VTA. Following recovery, all mice were initially tested for signal strength and responsivity of neural activity before inclusion in the study.
